# Hydrothermal pretreatment renders peat susceptible to enzymatic saccharification

**DOI:** 10.1101/2024.10.05.616816

**Authors:** Jonas Thomsen, Signe Lett, Helle Martens, Helle Sørensen, Darragh Kelleher, Theodora Tryfona, Paul Dupree, Katja S. Johansen

## Abstract

*Sphagnum* peat bogs store a large fraction of biologically-bound carbon, due to a steady accumulation of plant material over millennia. The resistance of *Sphagnum* biomass to decay is poorly understood but of high importance for preservation efforts and climate models. It is shown that peat cellulose and other glucose-rich polysaccharides are readily degradable by a commercial enzyme cocktail designed for the industrial saccharification of lignocellulose of vascular plants. However, prior hydrothermal pretreatment of peat was required for the enzymes to gain access to the polysaccharides. The pretreatment itself released monosaccharides and glucose-containing soluble oligosaccharides. The monosaccharide profile released from hydrothermal pretreatment was consistent with the expected hemicellulose content of *Sphagnum* and was clearly different from that seen for pretreated tissue from vascular plants, such as pretreated wheat straw. Cellulose retained in the insoluble part of peat was cleaved at a similar or higher rate compared to cellulose from vascular plant tissues. Confocal laser scanning microscopy showed that the hydrothermal pretreatment disrupted the cells and relocated lignin-like compounds. Peat contains a high concentration of iron, which likely explains the pronounced acidification observed for the pretreated peat slurry at ambient conditions during saccharification assays. The acidification is due to abiotic oxidative reactions that also inactivate the enzymes. Adding catalase to the reactions alleviated enzyme inactivation and essentially stopped acidification during saccharification. This study confirms the importance of considering those abiotic oxidative reactions that take place in drained peat material.

## 1. Introduction

Soil organic matter contains more organic carbon than the global vegetation and the atmosphere combined (Lehmann & Kleber, 2015). Of the estimated 4,000 billion tons of soil carbon, an estimated 500 billion tons is stored in northern peatlands (Yu, 2012). In light of the current climate crisis, the importance of keeping these vast carbon stores in place has led to international calls for peatland restoration and protective legislation (Environment programme. UNEP. 2024; Nature Restoration Law. European Commission. 2022). Peat profiles in bogs show increasingly decomposed plant material to a depth of several metre, representing thousands of years of carbon accumulation. Although the plant matter is more decomposed the deeper it is below the surface, the carbon stored in deep waterlogged anoxic zones (the catotelm) is very stable (Wilson et al., 2016). This remarkable recalcitrance and preservation of peat has been a matter of debate, with several hypotheses proposed to explain these properties. For example, the slow decomposition rate of the top layer (the acrotelm) has been attributed to increasingly anoxic conditions, the composition of the plant cell wall polysaccharides (Hájek et al., 2011), and the phenolic content of *Sphagnum*, the main vegetation comprising the top layer (Freeman et al., 2001). The low concentration of nutrients and acidic pH also likely contribute to the preservation of peat (Moore & Basiliko, 2006). More recently, it was proposed that soil organic matter in peat bogs is stabilized by complexing with iron (Y. Wang et al., 2017).

Vascular plants synthesize lignin to strengthen cell wall integrity and protect polysaccharides from degradation. Bryophytes, including *Sphagnum*, do not synthesize lignin but do contain polyphenolic lignin-like compounds (Weng & Chapple, 2010). These lignin-like compounds have been detected in Sphagnum leaves by histological approaches and from detecting autofluorescence intensities similar to that of vascular plants (Ligrone et al., 2008). Electron microscopy analysis of cryo-fractured hyaline leaf cells from *Sphagnum fuscum* suggested that the lignin-like compounds surround the cellulose, possibly forming a protective amorphous layer that could restrict access by cellulases (Tsuneda et al., 2001). However, the removal of phenolics had negligible effects on the degradation of *Sphagnum* polysaccharides by a microbial inoculum from a bog (Hájek et al., 2011). *Sphagnum* likely has important composition and cell wall architectural adaptations leading to greater recalcitrance compared to vascular plants. For example, fungal taxa isolated from a boreal peat bog decomposed spruce wood chips faster than *Sphagnum fuscum* biomass (Thormann et al., 2002). *Sphagnum* was thought to contain an unusual pectin-like polymer, sphagnan (Painter, 1983) with tanning properties (Painter, 1991), but this claim has since been disputed (Ballance et al., 2007). Overall, the reason behind the particular recalcitrance of *Sphagnum* to decomposition remains poorly understood.

Several environmental factors have been proposed to influence the preservation of peat. In the enzymatic latch hypothesis, putative *Sphagnum*-degrading enzymes are inhibited by phenolic compounds, with phenol oxidase as a key enzyme preventing this inhibition by degrading the inhibiting phenolic compounds (Freeman et al., 2001). Recent studies have challenged this hypothesis showing no or little relation between *Sphagnum* degrading enzymes and concentration of phenolics (Romanowicz et al., 2015; Urbanová & Hájek, 2021), including some that focus on the role of iron in peat preservation. The “iron gate” theory suggests that when oxygen enters the peat due to receding water levels, ferrous iron Fe(II) is oxidized to ferric iron Fe(III), which then complexes with phenolic compounds, thus creating an “iron gate” against phenol oxidase, protecting the phenolic compounds from degradation (Y. Wang et al., 2017). However, a study investigating the effect of drainage on the protection of soil carbon by metals in peatland dominated by *Sphagnum* or non-*Sphagnum,* reported that the “iron gate” mechanism is highly influenced by the vegetation type. In a *Sphagnum-*dominated peatland, most carbon was metallic bound under waterlogged conditions compared to drained conditions, while the opposite was true for non-*Sphagnum* peatland (Liu et al., 2024). The role of iron in the preservation of carbon in wetlands, particularly in *Sphagnum-*dominated wetlands, thus remains unclear.

The recalcitrance of plant cell walls is partially due to the high chemical stability of the 1-4 β-glycosidic bond. Indeed, the polysaccharides by themselves have a half-life on the order of millions of years (Wolfenden & Snider, 2001). Secreted microbial enzymes, which synergistically degrade cellulose and hemicellulose, induce turnover rates down to seconds, however. In vascular plants, limited access (at the nano-scale level) to the polysaccharides can be the main factor limiting such enzymatic degradation of lignocellulosic cell walls (Angeltveit et al., 2023; Johansen, 2016). Whether this also holds true for *Sphagnum* cell walls remains unknown. It is generally acknowledged that S*phagnum* contains cellulose (Kremer et al., 2004; Pipes & Yavitt, 2022), but the similarity of *Sphagnum* cellulose to that of vascular plants has not been fully clarified. Early work on the nature of cellulose in *Sphagnum,* extracted and chemically characterized cellulose from S*phagnum recurvum* and from cotton fibers. The cellulose of *Sphagnum* behaved differently from that of cotton during extraction, and the extracted cellulose exhibited chemical differences compared to cotton. Whether the limit to enzymatic degradation of *Sphagnum* cell walls is attributable to limited access to cellulose and other polysaccharides, or possibly to differences in cellulose, remains unknown.

The enzymatic digestion of lignocellulose releases saccharides and is therefore termed saccharification. This process have been optimised through decade-long research initiatives and is now applied in industrial settings for efficient conversion of lignocellulosic biomass, such as pretreated sugar cane bagasse, corn stover and wheat straw into ethanol (Johansen, 2016). When saccharification conditions such as temperature, pH, and dissolved oxygen content are controlled and optimised, an enzyme cocktail can degrade more than 80% of cellulose from steam-pretreated wheat straw (Scott et al., 2016). The degree to which peat would be degraded under such conditions is unknown. However, peat has long been used in horticulture as growth medium for soil improvement. Under these conditions, it is well known that peat decomposes. This leads to the main research question addressed in this study: How recalcitrant is the cellulose in peat to saccharification by an enzyme cocktail optimised for lignocellulose degradation?

In this study, we investigated the enzymatic digestibility of peat polysaccharides by applying methods developed for enzymatic degradation of lignocellulose from vascular plants. It is shown that peat is resistant to saccharification by an industrial enzyme cocktail. However, hydrothermal pretreatment of peat renders the material highly accessible to enzymatic saccharification. The overall catalytic rate and inactivation constants for the commercial enzyme cocktail used for saccharification of pretreated peat are estimated. Confocal laser scanning microscopy confirmed the disruption of cells and dislocation of cell wall components by the pretreatment. These findings suggests that access of the polysaccharide degrading enzymes to their substrates is a major reason for the recalcitrance of peat to degradation. Thus, this study provides new insight on the particular recalcitrance of *Sphagnum* peat.

## 2. Methods

### 2.1. Enzymes and reagents

All reagents were laboratory grade, unless otherwise stated. The commercial enzyme cocktail Cellic® CTec3, catalase, and lactrol for preventing growth of lactic acid bacteria, were kind gifts from Novonesis A/S.

### 2.2. Substrates

Non-fertilized garden peat (Silvan, Grønne Fingre), hereafter referred to as peat, was used as the main peat substrate in this study. Peat was sieved through a 1 mm mesh to remove stones and small pieces of wood and was then stored at room temperature in a closed plastic bucket. For comparisons, acrotelm and catotelm collected from a known mire (peatland) and hydrothermally pretreated wheat straw were used. Hydrothermally pretreated wheat straw is a well described lignocellulosic material. Steam-pretreated wheat straw was previously prepared by Kristensen et al., 2008 and stored at −18°C. The pretreated wheat straw was thawed at 4 °C prior to saccharification experiments. The wheat straw had been pretreated at 195 °C with a residence time in the pretreatment reactor of six minutes. Acrotelm and catotelm was were sampled from the palsa mire ‘Storflaket’ (68°20′48′′N,18°58′16′E) close to Abisko Scientific Research station in subarctic Sweden. The sampling spot was typical *Sphagnum fuscum* (Schimp.) H. Klinggr. hummock. Specifically, on July 22, 2023, a hole was cut with a bread knife to a depth of approximately 40 cm under which the peat was frozen. Acrotelm was collected from just below the capitula of the moss shoots and to 10 cm in depth. Catotelm peat was collected just above the frozen layer. Samples were transported in plastic bags on ice. All collected materials originated from *Sphagnum-*dominated bogs and therefore consisted mainly of *Sphagnum* cell wall remains.

### 2.3. Determination of substrate composition

The composition of structural polymers in the catotelm, untreated peat and pretreated peat was determined by sulphuric acid hydrolysis in triplicates. A wheat straw reference material from the National Institute of Standards and Technology (NIST) was analysed in parallel to validate the composition analysis. First, the plant materials were oven-dried at 50°C for two days. Afterwards, the samples were ground with a mortar and pestle and the dry matter (DM) content was determined in triplicate using a moisture analyzer (HC103, Metler Toledo). Approximately 15 mg dried material was weighed and placed into 2.5-mL screw cap tubes, soaked in 83 µL of 72% (v/v) sulphuric acid and incubated at room temperature for 1 hour with vortexing every 10 minutes. The volume was adjusted to 1.5 mL with deionized water, resulting in a 4% (v/v) sulphuric acid solution and 1% (w/v) biomass before incubation at 121 °C for one hour in a heating block. Finally, samples were cooled on ice and vacuum filtered through a pre-weighted sintered glass filter crucible por. 4 (Duran, Buch og Holm, Denmark) to separate the acid-soluble and acid-insoluble fraction. The acid-soluble fraction was neutralized with CaCO_3_ and spun for 10 minutes at 2000 rpm. The supernatant was filtered through sintered 0.45-µm glass filters, and analysed for arabinose, galactose, glucose, xylose and mannose on a Dionex ICS-5000 (Thermo Scientific). Monosaccharide content was converted into the theoretical amount of homopolymers as previously described (Santana & Okino, 2007). The acid-insoluble part of the biomass, Klason lignin + ash, was retained in glass the filter and measured gravimetrically after drying at 105 °C overnight.

### 2.4. Analysis of metal contents in the substrates

Transition metals are involved in oxidative chemistry. The concentration of metals including iron, copper, manganese, aluminum was quantified in the substrates by inductively coupled plasma-mass spectrometry (ICP-MS). The substrates were first freeze-dried and homogenized with a mortar and pestle. Then, 0.3 g of each substrate was weighed into a Teflon bomb and 10 mL of concentrated nitric acid (70%) was added. The samples were heated in a microwave with a ramping phase of 20 minutes and a hold phase of 25 minutes at 180 °C. After cooling, the solutions were first diluted to 50 mL with MilliQ water and further diluted 10-fold before analysis.

### 2.5. Pretreatment of peat

A slurry of 15% (w/w) dry matter was prepared in deionized water and had a pH of 4.11, which was not adjusted before pretreatment. The pretreatment at 121 °C was carried out in an autoclave and the pretreatments at 140°C, 160°C and 180 °C were performed in a Parr reactor (model 4848, PARR Instrument Company) with continuous stirring at 100 rpm and a 5 minute hold phase at the desired temperature (Fabiola Rodríguez-Zúñiga et al., 2015).

### 2.6. Saccharification with CTec3

Peat, pretreated peat substrates and pretreated wheat straw were enzymatically saccharified essentially as described previously for steam-pretreated wheat straw (Scott et al., 2016). Briefly, 20 g of a slurry of 5% (w/w) DM was hydrolysed in duplicate samples in 50 ml falcon tubes at 50 °C and pH 5.2 in a hybridization incubator (combi-D24, Finepcr) for up to 144 hours. Lactrol was added to a concentration of 75 mg/kg slurry to prevent growth of lactic acid bacteria. The commercial enzyme cocktail CTec3 was used in three dosages of 5, 10, and 15 mg/ g DM. In addition, the effect of 10, 50 and 100 µL catalase on saccharification efficiency of pretreated peat incubated with 15 mg CTec3 was assessed. To determine the glucose concentration during saccharification, samples were taken from the falcon tubes after 1.5, 3, 6, 24, 48 and 72 hours of incubation. After each sampling, the pH was adjusted with 1 M KOH, except at the 3 hours sampling time. The concentration of monosaccharides was determined by high-performance liquid chromatography (HPLC) analysis using a Dionex ICS-5000 as described below. Fractional conversion of cellulose was calculated by dividing the glucose concentrations, measured by HPLC at each time point, by the original cellulose content of the substrate. The soluble glucose concentration prior to enzyme addition and the amount of glucose added with CTec3 was subtracted from the measured glucose before calculating the fractional conversion.

### 2.7. Washing of peat pretreated at 180 °C

To separate soluble and insoluble glucan in peat pretreated at 180 °C, a slurry of 1% (w/w) DM was prepared in a 250-mL plastic bottle. The 1% DM slurry was shaken for 10 seconds by hand and then centrifuged at room temperature for 5 minutes at 3000 rpm. The supernatant was analysed for monosaccharides and soluble oligosaccharides by HPLC. This procedure was repeated 10 times, and the remaining insoluble part of the pretreated peat was used in saccharification experiments.

### 2.8. Saccharification of soluble glucan released from peat pretreated at 180 °C

The soluble part in the first wash of peat pretreated at 180 °C was centrifuged at 14,000 rpm for 10 minutes at room temperature and sterile-filtered (0.45 µm). Duplicate samples of 1 mL were incubated for 1 h at 50 °C in a thermomixer (Eppendorf) with shaking at 600 rpm with 5 µL of CTec3 diluted 10-fold from the commercial product added to each sample. The samples with and without CTec3 were analyzed for their contents of monosaccharides and oligosaccharides by HPLC as described below. Any content of monosaccharides or oligosaccharides in CTec3 was below the HPLC detection limit.

### 2.9. HPLC analysis of monosaccharides, oligosaccharides and acetate

The contents of monosaccharides (arabinose, galactose, glucose, xylose, mannose) from saccharification experiments, compositional analysis and pretreatment assessment, as well as oligosaccharides, were analyzed by high-performance anion-exchange chromatography (HPAEC) using a Dionex ICS-5000 (Thermo Scientific) equipped with a CarboPac PA1 column (Thermo Scientific) and a Dionex IonPac guard column (Thermo Scientific) operated at 30 °C and with pulse amperometric detection (PAD). Monosaccharides were eluted with MilliQ water as eluent at a flow rate of 0.25 mL/min and an isocratic flow of 0.2 M NaOH at a flow rate of 0.1 mL/min. Oligosaccharides were separated by applying a gradient 1 M sodium acetate as described by Westereng et al., 2013.

Samples of peat pretreated at 180 °C taken after 1.5, 3, 6, 24, 48 and 72 hours of incubation with and without CTec3 were also analyzed for acetate contents using a Dionex Ultimate 3000 system (Thermo Scientific) with a Rezex refractive index detector and a ROA-organic acid H+(8 %), 300 × 7.8 mm column (Phenomenex). The system was operated at 80 °C with 5 mM H_2_SO_4_ as eluent at a flow rate of 0.6 mL/min. The lower detection limit of acetate for this method was 0.2 g/L.

### 2.10. Polysaccharide analysis by carbohydrate gel electrophoresis (PACE)

Soluble part of peat pretreated at 180 °C and cello-oligosaccharide standards were derivatised with 8-aminonaphthalene-1,3,6-trisulfonic acid (ANTS) by reductive amination according to previous protocols. Derivatised samples were dried under vacuum and solubilised in 3M urea. For carbohydrate electrophoresis, samples were loaded on a 10% (w/v) polyacrylamide gel and electrophoresed at 10 °C at 1000 V for 1 h using 0.1 M Tris-borate (pH 8.2) as running buffer. A GBox CCD camera with a transilluminator with long-wave tubes emitting at 365 nm was used for PACE gel scanning. Images were captured using GENESNAP software.

### 2.11. Enzyme kinetics

Building on previous work (Peciulyte et al., 2018; Scott et al., 2016), a two-stage kinetic model for enzymatic release of glucose was used to estimate the inactivation *k_i_* and the catalytic *k_s_* rate constants, taking into account the conversion of cellulose to cellobiose (first stage) and cellobiose to glucose (second stage) by a complex enzyme cocktail. The model consists of a set of five differential equations that include enzyme specific constants (Scott et al., 2016). The input for the equations are the concentration of cellulose (S), cellobiose (G2), glucose (G), and active cellulase (E_a_). Optimal values of *k_i_* and *k_s_* were determined by non-linear least squares, keeping other parameters fixed at the same values given in previous work (Peciulyte et al., 2018; Scott et al., 2016). Computations were carried out with custom-made code in R (version 4.4.1) (*R: The R Project for Statistical Computing*) using the deSolve package (version 1.35) (Soetaert et al., 2010) to numerically solve the system of differential equations with the Dormand-Price method and the optim function to minimize the least square’s function (Appendix B). Previous work with the kinetic model used Excel Solver for the computations (Scott et al. 2016). The model was applied to data from 180 °C pretreated peat and the insoluble part of 180 °C pretreated peat. To validate the R code, it was applied to the fractional conversion data of pretreated wheat straw from this study, giving *k*_i_ and *k*_s_ values very similar to those calculated in Scott et al. 2016 (Fig. S1A).

### 2.12. Microscopy

Leaf material from frozen samples of *Sphagnum fuscum* plants was thawed and immediately fixed in Karnovsky’s fixative (5% [w/v] glutaraldehyde, 4% [w/v] paraformaldehyde, 0.1 M sodium cacodylate buffer, pH 7.3), and stored at 4°C until analysed, together with fixed samples of peat and 180 °C pretreated peat. All preparations were performed in triplicates. Leaf tips were stained for 15 minutes in 0.5% (w/v) calcofluor white, rinsed in distilled water, and mounted onto glass slides in a 20 μL droplet of distilled water and imaged with a Leica SP5X confocal laser scanning microscope (Mannheim, Germany) using a 20×objective. Calcofluor white was excited with the UV laser (355 nm), and emission was collected at 391–463 nm. An excitation wavelength of 488 nm and an emission band of 513–556 nm was used for the detection of autofluorescence from secondary cell walls.

## 3. Results and discussion

To investigate the extent to which *Sphagnum* peat is susceptible to enzymatic saccharification, we used an approach similar to that applied to lignocellulosic materials targeted for industrial bioconversion. Specifically, we conducted a chemical characterisation, hydrothermal pretreatment, assessment of abiotic reactivity, and enzymatic saccharification. Furthermore, we used an enzyme kinetic model to calculate the catalytic rate and first-order inactivation rate constants of the cellulases present in CTec3, the enzyme cocktail used for enzymatic saccharification Garden peat (peat) was used as the primary source of peat for this study and compared to those obtained for environmental samples from a peat bog (catotelm and acrotelm). We also used steam-pretreated wheat straw as a well described and industrially relevant lignocellulosic material for comparison.

### 3.1. Chemical composition of peat substrates and pretreated wheat straw

The composition of the main constituents of lignocellulose, namely cellulose, hemicellulose and the residual fraction of Klason lignin and ash, were determined by sulphuric acid hydrolysis (Table 1). While *Sphagnum* does not contain lignin *per se*, the peat samples had a much higher content of Klason lignin and ash than steam-pretreated wheat straw. The content of cellulose was calculated based on the glucose concentration in the sulphuric acid–hydrolysed samples. This method is commonly used for a subsequent assessment of biomass saccharification (Merali et al., 2015; Petersen et al., 2009). However, this method is not entirely accurate, as glucose will also be released from polysaccharides other than cellulose during sulphuric acid hydrolysis. Nevertheless, we estimated the cellulose content to be 4–5 times lower in peat (128 mg/g) and catotelm (114 mg/g) than in steam-pretreated wheat straw (567 mg/g). The content of hemicellulose was calculated based on the concentration of arabinose, galactose, xylose and mannose in the acid-hydrolysed samples, which reached levels roughly twice as high in the *Sphagnum* samples as in steam-pretreated wheat straw. Because different methods for determining the composition of peat have previously been used (Pipes & Yavitt, 2022; Xu et al., 2021), it is not straightforward to directly compare our results and those from other studies.

**Table 1.**
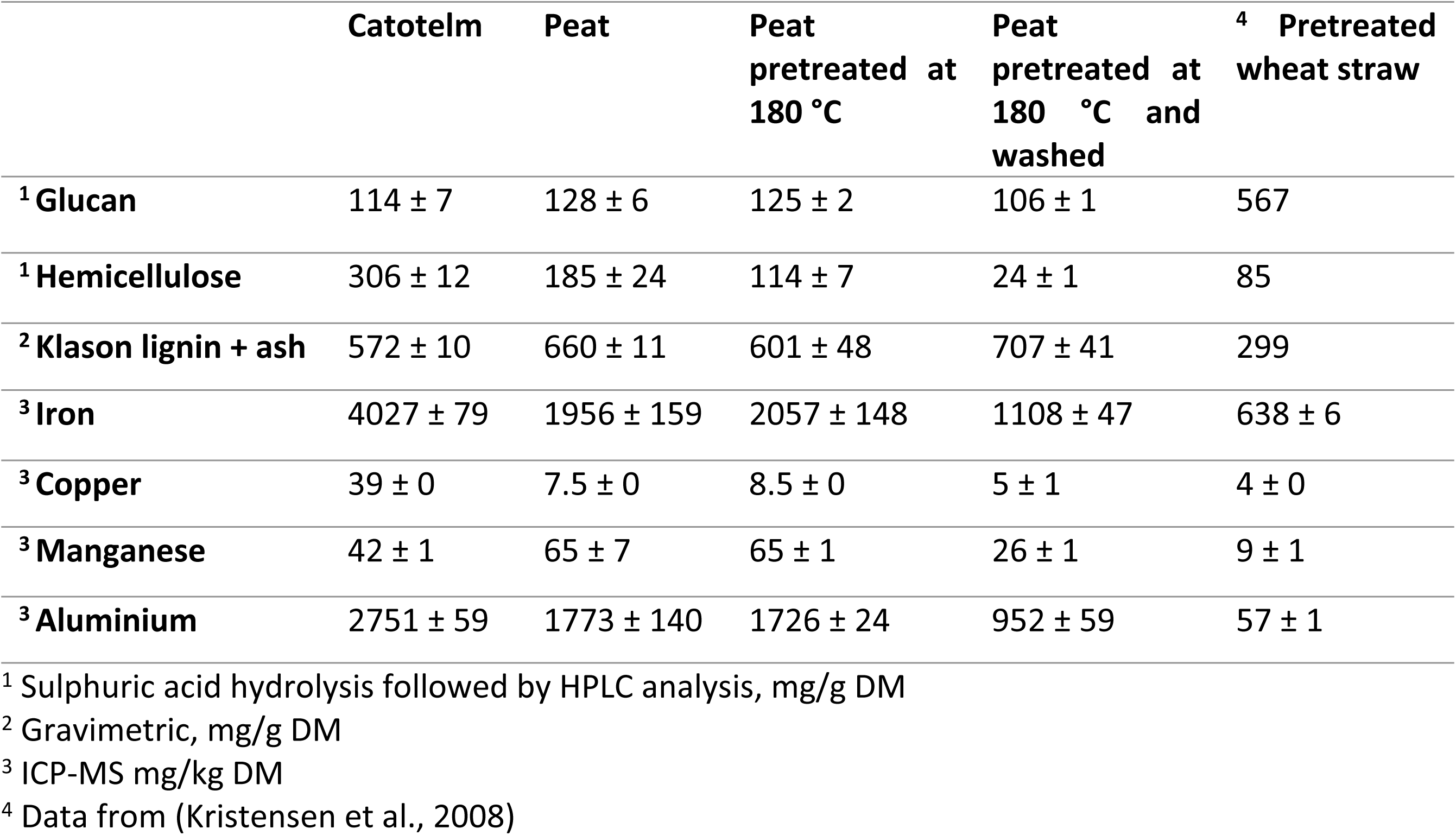
Summary of composition of structural polymers and concentrations of common redox-related metals in catotelm peat, peat samples and pretreated wheat straw. Values are presented as means ± standard deviation (SD) from three biological replicates. Pretreatment of peat is described in section 3.2.

|  | Catotelm | Peat | Peat<br>pretreated at<br>180 °C | Peat<br>pretreated at<br>180 °C and<br>washed | <sup>4</sup> Pretreated<br>wheat straw |
| --- | --- | --- | --- | --- | --- |
| <sup>1</sup> Glucan | 114 ± 7 | 128 ± 6 | 125 ± 2 | 106 ± 1 | 567 |
| <sup>1</sup> Hemicellulose | 306 ± 12 | 185 ± 24 | 114 ± 7 | 24 ± 1 | 85 |
| <sup>2</sup> Klason lignin + ash | 572 ± 10 | 660 ± 11 | 601 ± 48 | 707 ± 41 | 299 |
| <sup>3</sup> Iron | 4027 ± 79 | 1956 ± 159 | 2057 ± 148 | 1108 ± 47 | 638 ± 6 |
| <sup>3</sup> Copper | 39 ± 0 | 7.5 ± 0 | 8.5 ± 0 | 5 ± 1 | 4 ± 0 |
| <sup>3</sup> Manganese | 42 ± 1 | 65 ± 7 | 65 ± 1 | 26 ± 1 | 9 ± 1 |
| <sup>3</sup> Aluminium | 2751 ± 59 | 1773 ± 140 | 1726 ± 24 | 952 ± 59 | 57 ± 1 |

<sup>1</sup> Sulphuric acid hydrolysis followed by HPLC analysis, mg/g DM
<sup>2</sup> Gravimetric, mg/g DM
<sup>3</sup> ICP-MS mg/kg DM
<sup>4</sup> Data from (Kristensen et al., 2008)

Oxidative abiotic reactions can promote or impede the degradation of lignocellulose and involve transition metals such as iron and copper (Miles & Brezonik, 1981; Peciulyte et al., 2018). Significant decarboxylation (CO_2_ production) of organic matter in natural humic-coloured waters, catalysed by iron, has been demonstrated (Miles & Brezonik, 1981). Such abiotic reactions are also taking place in slurries of pretreated wheat straw under saccharification conditions (pH 5.3 and 50 °C), involving H_2_O_2_ as an intermediate. The resulting CO_2_ production contributes to acidification of the saccharification slurry (Peciulyte et al., 2018). We therefore carried out an ICP-MS analysis to determine the metal content of all samples. High concentrations of iron was observed in the peat samples (Table 1, Fig. S2).

In particular, the catotelm contained 4000 mg of iron per kg of dry matter (DM), while peat contained 2000 mg iron per kg DM, which was 6-fold and 3-fold higher than in the wheat straw used in this study, respectively. The high concentrations of iron and aluminum in the catotelm and peat are consistent with reports that peatlands globally tend to store large quantities of metals, with nutrient-poor bogs containing 2500 mg iron per kg on average and nutrient-rich fens up to 16.000 mg iron per kg (Osborne et al., 2024). *Sphagnum*-derived peat, in particular, has been suggested to be prone to accumulating metals because of complexation with acidic and phenolic compounds from *Sphagnum* cell walls (Clymo, 1963; Riedel et al., 2013; Zhao et al., 2023). From the degradation of peat, a substantial pool of iron might therefore be released and lead to a noticeable degree of abiotic oxidative reactions.

### 3.2. Hydrothermal pretreatment of peat and its effect on acidification of peat slurries

We hypothesised that hydrothermal pretreatment could enable enzymatic saccharification of peat. Therefore, a slurry of 15% (w/w) DM in water was pretreated at either 121 °C by autoclaving or at 140°C, 160°C, or 180°C in a Parr reactor. Due to the high iron content in peat samples (Table 1), it was necessary to evaluate to which extent the peat slurries would acidify, under the same conditions as for the subsequent saccharification (50 °C, pH 5.2). Peat pretreated at high temperatures (160 °C or 180 °C) required a higher volume of 1 M KOH to maintain the slurry pH to 5.2 (85 µL and 100 µL, respectively), than for peat samples pretreated at the lower temperatures (40 µL for 120 °C and 45 µL for 140 °C). This finding suggested a strong correlation between slurry acidification and pretreatment temperature (Fig. 1A). Disruption of chemical bonds by the pretreatment likely led to release of iron otherwise sequestered by negatively charged compounds, thereby increasing the production of CO_2_. Uronic acids like galacturonic acid may be released by the hydrothermal pretreatment of peat and contribute to acidification, but decompose into their degradation products like furfurals at high temperatures (Benito-Román et al., 2024; Pińkowska et al., 2019). Release of acetic acid from hemicelluloses also contributes to acidification during the saccharification of lignocellulose slurries (Peciulyte et al., 2018). However, the concentration of acetic acid in the pretreated peat slurries was below the detection limit (data not shown).

**Fig. 1.**
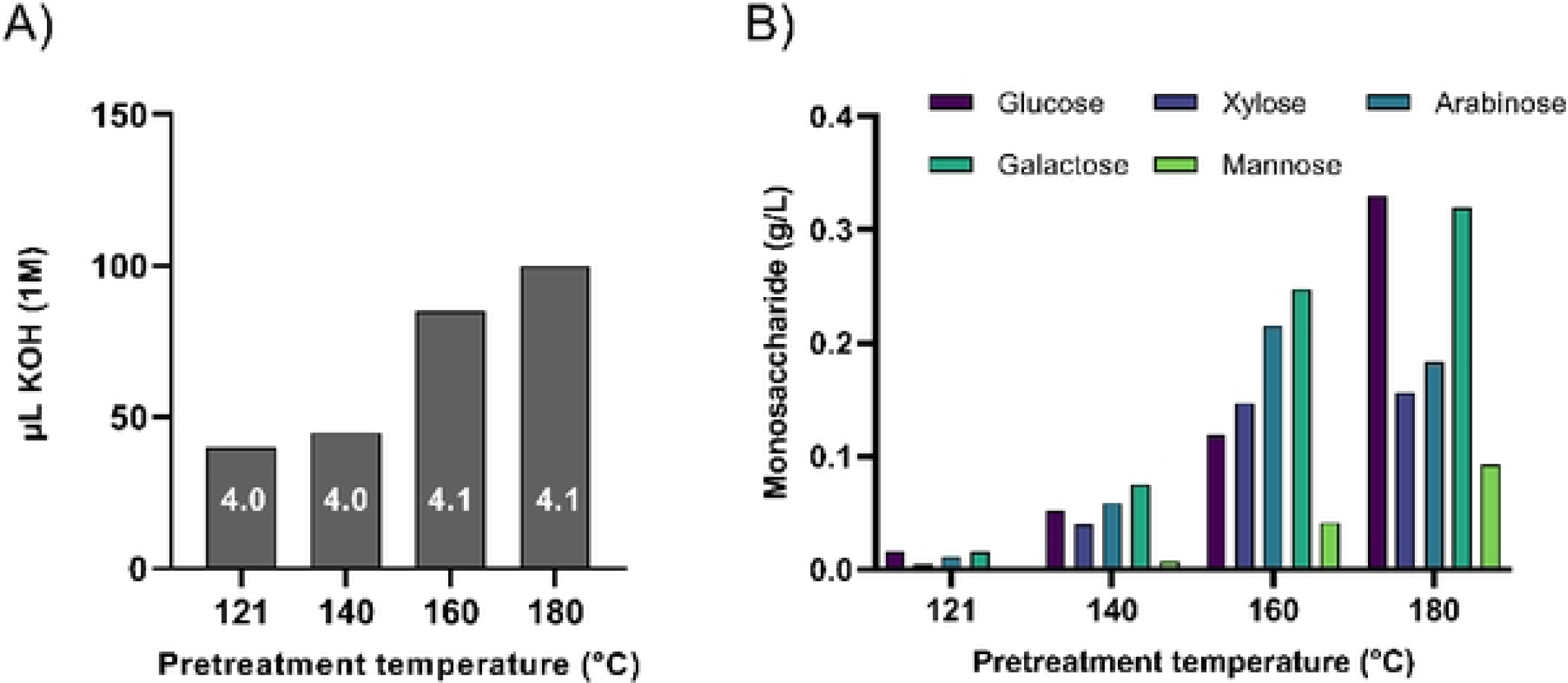
Acidification and monosaccharide content of pretreated peat. A) Volume of 1 M KOH required to keep the pH at 5.2 during 24 h of incubation without CTec3 at 50 ’C. During incubation, pH was adjusted to 5.2 with 1 M KOH after 1.5, 6 and 24 hours. The same amount of KOH (1 M) was added to both duplicate samples of the 20 g slurry. The pH of each pretreated material without any adjustments is noted in each column. B) Monosaccharide solubilization from peat as a function of pretreatment temperature. The values are averages of duplicate samples.

The effect of pretreatment temperature on slurry acidification during incubation was therefore likely explained by an increase in abiotic oxidative reactions. As the concentration of dissolved oxygen under lab conditions is much higher than concentrations found in a peat bog, the effect on acidification observed here would not be expected to be as prominent below the water table in an undisturbed peat bog. However, such iron-catalysed reactions involving H_2_O_2_ as an intermediate are well described in environmental systems such as arctic soil waters (Page et al., 2013; Trusiak et al., 2018). It was also been shown that CO_2_ can be a main contributor to peatland water pH (Shotyk, 1988). In the uppermost part of peatlands where *Sphagnum* grows, organic acids such as uronic acids and phenolic acids released from *Sphagnum* cell walls also contribute to an acidic pH (van Breemen, 1995; Verhoeven & Liefveld, 1997).

### 3.3. Hydrothermal pretreatment of peat releases monosaccharides

Hydrothermal pretreatment of plant materials partially disrupts chemical bonds in the cell wall (Ibbett et al., 2014). Depending on the pretreatment conditions, hemicelluloses such as xylan are degraded to a greater or lesser extent, thus releasing monosaccharides. These physicochemically released monosaccharides must be quantified before the enzymatic saccharification efficiency can be assessed. The concentration of monosaccharides in the soluble part increased with pretreatment temperature, as expected (Fig. 1B). While hardly any monosaccharides was released from pretreatment at 121 °C, all five monosaccharides were in concentrations from about 0.1–0.3 g/L in the soluble part of peat pretreated at 180 °C. Noticeably, the glucose concentration increased from 0.12 g/L with the pretreatment temperature of 160°C to 0.33 g/L with the 180°C pretreatment. The concentrations of the other sugars in the 180°C pretreated sample were 0.32 g/L for galactose, 0.18 g/L for arabinose, 0.16 g/L for xylose, and 0.09 g/L for mannose.

The polysaccharide composition of peat is largely unknown. This material may contain plant residue from more than one species that has been partially degraded during hundreds of years in the bog and during excavation and storage. However, the monosaccharide profile released by the 180 °C pretreatment, with high concentrations of glucose, galactose and arabinose, is very different from the typical xylan-rich monosaccharide profile of wheat straw pretreated at 180 °C (Thygesen et al., 2004). This observation suggested major differences in cell wall architecture between the two types of plants. The monosaccharides in the pretreated peat are likely constituents of *Sphagnum* cell wall polysaccharides such as arabinoglucan, xyloglucan and glucomannan (Ye & Zhong, 2022). Arabinoglucan is a moss-specific polysaccharide first identified in *Physcomitrella patens* and is partially soluble when samples are heated to 60 °C in water (Roberts et al., 2018). Arabinoglucan is an unbranched and unsubstituted polysaccharide consisting of (1,4)-glucose and (1,3)-arabinose in a 7.5:1 ratio. Such a polysaccharide would very likely decompose into monosaccharides to a much larger extend than cellulose during hydrothermal pretreatment. The concentration of arabinose decreased from 160 °C to 180 °C, reflecting its degradation at high temperature (Sun et al., 2015; Zakaria et al., 2015). The relatively high concentrations of galactose measured here are supported by previous findings showing significant antibody labeling of galactan in hyaline cell walls of *Sphagnum* leaves (Kremer et al., 2004) and in the pectin-like polysaccharide sphagnan, which is extractable from the *Sphagnum* cell wall by heating to 98 °C in water (Ballance et al., 2007). Mannans are abundant hemicelluloses in the cell wall of several moss species (Ye & Zhong, 2022). The fact that we detected mannose in the soluble part of pretreated peat is interesting because mannans from vascular plants have been reported to primarily degrade into oligosaccharides by hydrothermal pretreatment between 160 °C and 190 °C (Z.-W. Wang et al., 2016).

### 3.4. Pretreatment renders peat enzymatically degradable

Slurries of peat and pretreated peat were saccharified using the cellulase-rich enzyme cocktail CTec3 at three different dosages, using pretreated wheat straw as a positive control for saccharification by CTec3 (Fig. 2). The CTec3 cocktail contains enzymes that release acetic acid from acetylated hemicelluloses in pretreated wheat straw, contributing to the acidification of the slurry (Peciulyte et al., 2018). There was a clear dosage-dependent acidification of pretreated wheat straw during saccharification with CTec3 (Fig. S3). The pretreated peat samples also acidified during saccharification and required the addition of more KOH to maintain the pH at 5.2 than wheat straw. However, there was no apparent effect of enzyme dosage on pH and acetic acid was not detected in samples. The cause of acidification of pretreated peat slurry during saccharification is therefore likely due to the abiotic oxidative reactions catalysed by iron as discussed above.

**Fig. 2.**
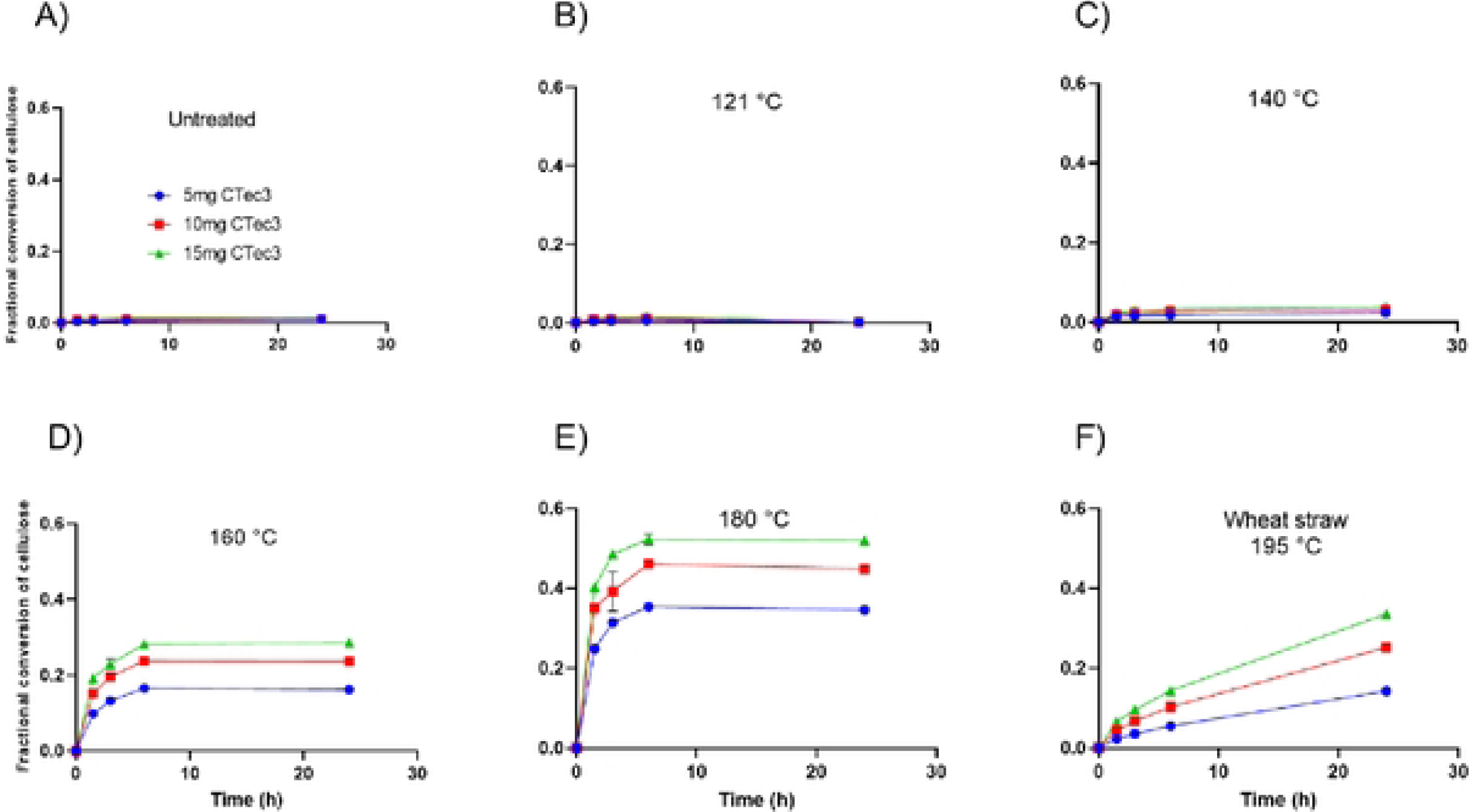
Effect of pretreatment temperature on peat saccharification progress curves. Fractional conversion of cellulose from A) untreated peat, B) peat pretreated at 121 °C, C) 140 °C, D) 160 °C, E) 180 °C, and F) pretreated wheat straw. All samples were incubated for 24 hours with 5 mg CTec3/g OM (blue), 10 mg/g OM (red) or 15 mg CTec3/g OM (green) at 50°C, at pH 5.2. The data are presented as means± SD from duplicate samples.

While minimal enzymatic conversion was observed for the untreated and autoclaved peat samples, a temperature-dependent increase in saccharification efficiency was seen for peat samples subjected to pretreatment at 140 °C, 160 °C, and 180 °C (Fig. 2). This enhancing effect of pretreatment on saccharification efficiency was similar to that previously seen for lignocellulose (Hansen et al., 2011). This result demonstrated that peat is similar to other types of plant biomass, in the sense that it requires a pretreatment step before enzymes can access their substrates in the plant biomass. The fractional conversion of cellulose, which is the degree to which the cellulose is converted to glucose, depended on enzyme dosage, reaching a maximum of 0.52 when using 15 mg CTec3/g DM from peat pretreated at 180 °C after 24 hour of incubation. The peat conversion progress curves showed a fast initial phase, followed by a rapid decline in further saccharification (Fig. 2E). These curves contrasted with the slow but steady conversion observed for pretreated wheat straw (Fig. 2F) that continued for 6 days (Fig. S1A). Since all conversion progress curves for pretreated peat samples stagnated after 6 hours, the enzymes were most likely inactivated rather than depleted of substrate. Both oxidative inactivation and substrate depletion were investigated further below.

### 3.5. Catalase reduce acidification and increases fractional conversion of pretreated peat

To investigate if the apparent inactivation of the enzyme cocktail was caused by oxidative reactions, we added catalase to scavenge the reaction intermediate H_2_O_2_. A previous examination of oxidative inactivation of cellulases during saccharification of wheat straw (Scott et al., 2016) indicated that the presence of oxidative enzymes called lytic polysaccharide monooxygenases (LPMOs) in the CTec3 cocktail decreased the half-life of cellulases, despite significantly increasing the glucose yield. This negative effect was however alleviated by addition of catalase, demonstrating that H_2_O_2_ is involved in the inactivation of the cellulases. Catalase was also shown to substantially decrease the requirements for KOH to maintain a constant pH, although there was no direct coupling between titrant requirements and sugar yield in that study (Peciulyte et al., 2018). Therefore, saccharification of peat pretreated at 180 °C was repeated with different dosages of catalase added.

First, the effect of catalase on titrant requirements was confirmed for the peat slurry, as all dosages of catalase tested clearly reduced the volume of KOH needed to maintain a pH of 5.2 during enzymatic saccharification of the peat slurry (Fig 3A). Without catalase, 165 µL KOH was required and with 10 µL catalase, only 50 µL KOH was required. Increasing the dosage of catalase to 100 µL led to a stable pH of in the peat slurry and no KOH was required during the 48 hours of incubation. Decreasing the amount of dissolved oxygen during saccharification of the GP slurry to around 1% (where 100% is air saturated) also lead to a decrease in KOH requirement substantially over 48 hours of saccharification (Fig. S4).

**Fig. 3.**
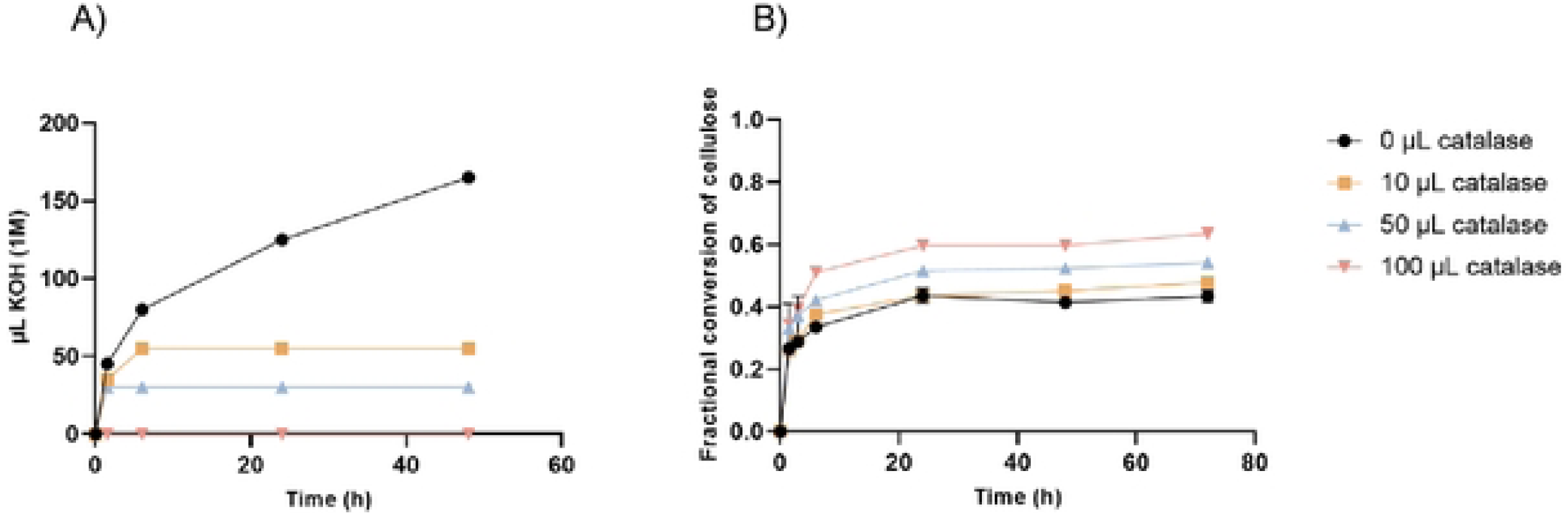
Effect of catalase on saccharification of 180°c pretreated peat. Duplicate samples of peat pretreated at 180°c were incubated for 72 hours with 15 mgCTec3 and a varying dosage of 0, 10, 50 or 100 µL catalase, at 50 °c and pH 5.2. A) Mean required amount of 1 M KOH to maintain the pH at 5.2 during saccharification. B) Mean fractional conversion of cellulose with varying dosages of catalase added.

Second, catalase increased saccharification of pretreated peat during the first 6 hours of incubation in a dosage-dependent manner, while the conversion progress curves stagnated for all treatments. This result contrasts with the positive effect of a 10 µL catalase addition on the saccharification of wheat straw which was most pronounced after 24 hours of incubation (Scott et al., 2016). The effect of catalase on enzyme inactivation during saccharification of peat slurries and the overall catalytic rate is discussed below.

### 3.6. Catalytic rate constant and inactivation constant for the saccharification of pretreated peat and pretreated wheat straw

To shed further light on the enzymatic saccharification of 180 °C pretreated peat, the two key kinetic constants for this heterogeneous and complex process was calculated. Previous work (Peciulyte et al., 2018; Scott et al., 2016) has presented a kinetic model describing the synergistic release of glucose from cellulose by a complex cocktail of enzymes and is used to estimate the inactivation rate constant (*k*_i_) and the catalytic rate constant (*k*_s_). This kinetic model consists of two stages, the first encompassing the conversion of cellulose to cellobiose by multiple enzymes and the second the production of glucose by β-glucosidase. However, when we applied the model to the fractional conversion data obtained from peat pretreated at 180 °C, the kinetic model exhibited a poor fit, suggesting that the fast initial phase of conversion followed by a rapid decline could not be explained by a combination of high catalytic rate and fast inactivation of the enzymes. A plausible alternative explanation would be that pretreated peat contains a considerable amount of soluble glucose-rich oligosaccharides or polysaccharides, which are highly accessible to the enzymes.

### 3.7. Pretreated peat contains both soluble and insoluble glucan

Overall, the shapes of the progress curves for saccharification of peat were clearly different from those obtained for pretreated wheat straw, and the initial rate of glucan conversion of peat pretreated at 180 °C was noticeably high. To test whether peat pretreated at 180°C contained a highly enzyme accessible soluble glucan, we extensively washed the peat slurry and collected the soluble and insoluble parts. The monosaccharide content in the soluble part after incubation with and without CTec3 was determined by HPLC, which showed that glucose is predominantly released by the enzyme cocktail (Fig. 4A). A range of oligosaccharides with retention times similar to those of cello-oligomers (1-4 β-linked glucose) with a degree of polymerization of 2–5 were detected in the soluble part (Fig. 4B). It is conceivable that cellooligomers could be released from cellulose to some extent, but hydrothermal pretreatment of wheat straw under conditions similar to those used in this study did not yield detectable amounts of cello-oligomers (Sidiras et al., 2011). Consistent with Sidiras et al., 2011, addition of H_2_SO_4_ during pretreatment of wheat straw at 180 °C leads to a very limited release of soluble cello-oligomers (Kabel et al., 2007). This suggests that either cellulose in peat is more extensively degraded during pretreatment or that a glucose-rich non-cellulosic polysaccharide is present in the soluble part. This is a possible outcome, as the oligosaccharide co-eluting with the cellotriose standard was not accessible to the CTec3 cocktail (Fig. 4C). Additionally, a range of oligosaccharides of lower abundance eluting between 13 and 23 minutes were resistant to the CTec3 cocktail. Polysaccharide analysis using carbohydrate gel electrophoresis (PACE) of the CTec3 treated soluble part indicated that the oligosaccharides are not derived from cellulose, since they did not co-migrate with cello-oligosaccharide standards (Fig. S5). Based on our estimates, the glucan in the soluble part of the pretreated peat accounted for 11% of the total glucan content (Fig. S6).

**Fig. 4.**
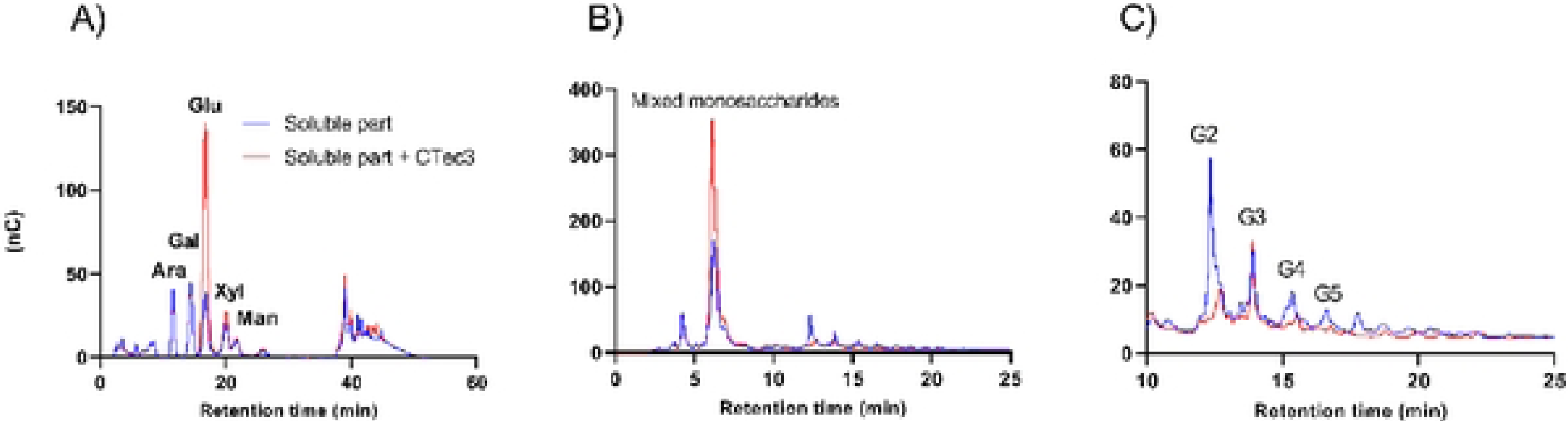
Release of glucan from soluble part after 1 hour incubation with or without CTec3 at 50 °c. A) Monosaccharide analysis of arabinose (Ara), galactose (Gal), glucose (Glu), xylose (Xyl), and mannose (Man). B) Oligosaccharide analysis of the soluble part with or without CTec3 treatment. C) Zoomed-in view of the peaks with retention times between 10 and 25 min from figure 4B. The peaks indicated G2-G5 have similar retention times as cellooligomers. Cellobiose (G2), triose (G3), tetraose (G4), pentose (GS).

### 3.8. Catalytic rate constant and inactivation constant for the saccharification of the insoluble part of peat

As the kinetic model was developed for the synergistic release of glucose from (insoluble) cellulose, it is arguably not applicable to produce kinetic constants from unseparated pretreated peat (Scott et al., 2016). Saccharification with CTec3 was therefore repeated with the insoluble part of the pretreated peat. The kinetic model was then applied and the new curves from the insoluble part of peat were a better, albeit not good, fit compared to those obtained for the non-washed peat (Fig. 5A). The catalytic rate constant *k*_s_ was calculated to 26.98 h^−1^ (SE ±2.61), which is two-fold higher compared to that of wheat straw. It is conceivable that the pretreatment led to more accessible chains on the outer part of peat cellulose, thus producing a higher *k*_s_ value. However, it is also possible that the pretreated peat contains other insoluble glucan containing polysaccharides.

**Fig. 5.**
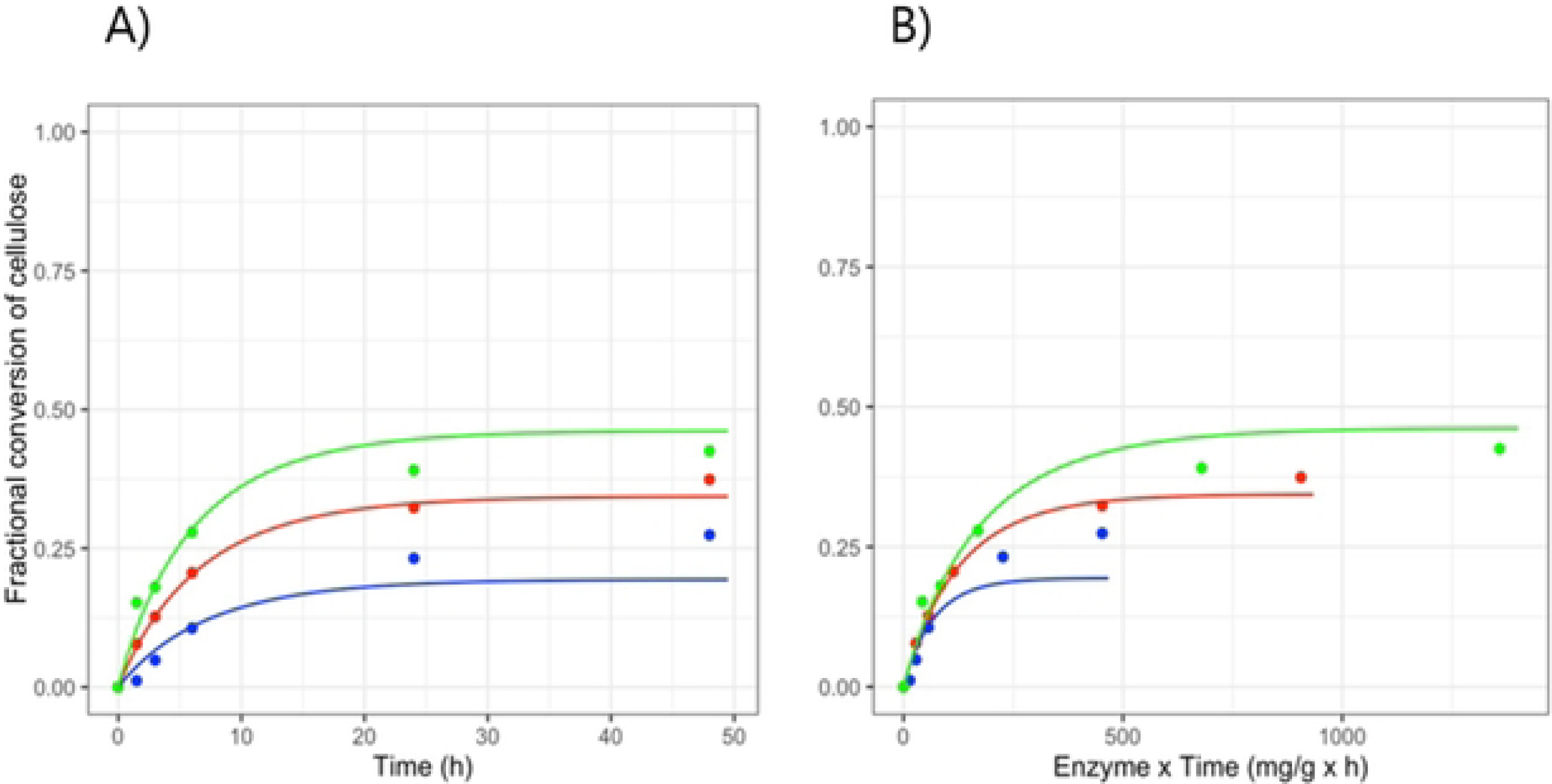
Inactivation of enzymes during saccharification of the insoluble part of pretreated peat. A) Fractional conversion of cellulose from the insoluble part of peat (washed 10 times) incubated with 5 (blue), 10 (red) and or 15 (green) mg CTec3/g OM for 48 hours at 50 °c, pH 5.2. B) To illustrate time­ dependent enzyme inactivation, the fractional conversion data was plotted as a function of enzyme dosage x time (mg/g x h).

The inhibition constant *k*_i_ was 125.91 ×10^−3^ h^−1^ (SE ± 15), thus around 6-fold higher than that for wheat straw (Fig. S1A). A higher *k*_i_ value means that the cellulolytic cocktail has a short half-life. The inactivation was also clear from a plot of the fractional conversion curves as a function of enzyme dosage x time (mg/g x h) (Fig. 5B) where curves lying on top of each other would illustrate no time-dependent inactivation (Selwyn, 1965). It is important to mention that the enzyme dosage for wheat straw and pretreated peat was the same when calculated based on DM, but the dosage of enzyme relative to cellulose content was 4.5 times higher for peat than for wheat straw due to the higher glucan content of wheat straw. Enzymes are in general stabilized by their substrates.

### 3.9. Visualization of changes in peat cell wall after pretreatment

Subjecting biomass to pretreatment will affect the cell wall integrity to some extent (Herbaut et al., 2018). *Sphagnum* has a unique leaf anatomy compared to other mosses and vascular plants, as they consist of dead and empty hyaline cells and chloroplast-containing chlorophyllose cells (*Glime, J. M. 2017.*). The effect of hydrothermal pretreatment on cell wall integrity of peat was investigated by confocal laser scanning microscopy. Calcofluor white was used to stain β-1,4 glucans such as cellulose, and lignin-like compounds were detected by autofluorescence. *Sphagnum* leaves from acrotelm were compared with the partially degraded catotelm. The acrotelm (Fig. 6A) showed the expected regularly arranged hyaline cells and chlorophyllose cells. The secondary cell walls of the hyaline cells strongly fluoresced in yellow/orange corresponding to lignin-like compounds, while the narrow chlorophyllose cells had clear cellulose-rich primary cell walls, with the narrow chlorophyllose cells having clear cellulose-rich primary cell walls. Thus, cellulose appears to be surrounded by a lignin-like compound, a similar observation as (Tsuneda et al., 2001). The autofluorescence signal from catotelm (Fig. 6B), collected about 40 cm below the acrotelm, was significantly lower than that of the acrotelm, but with some degree of cell integrity left.

**Fig. 6.**
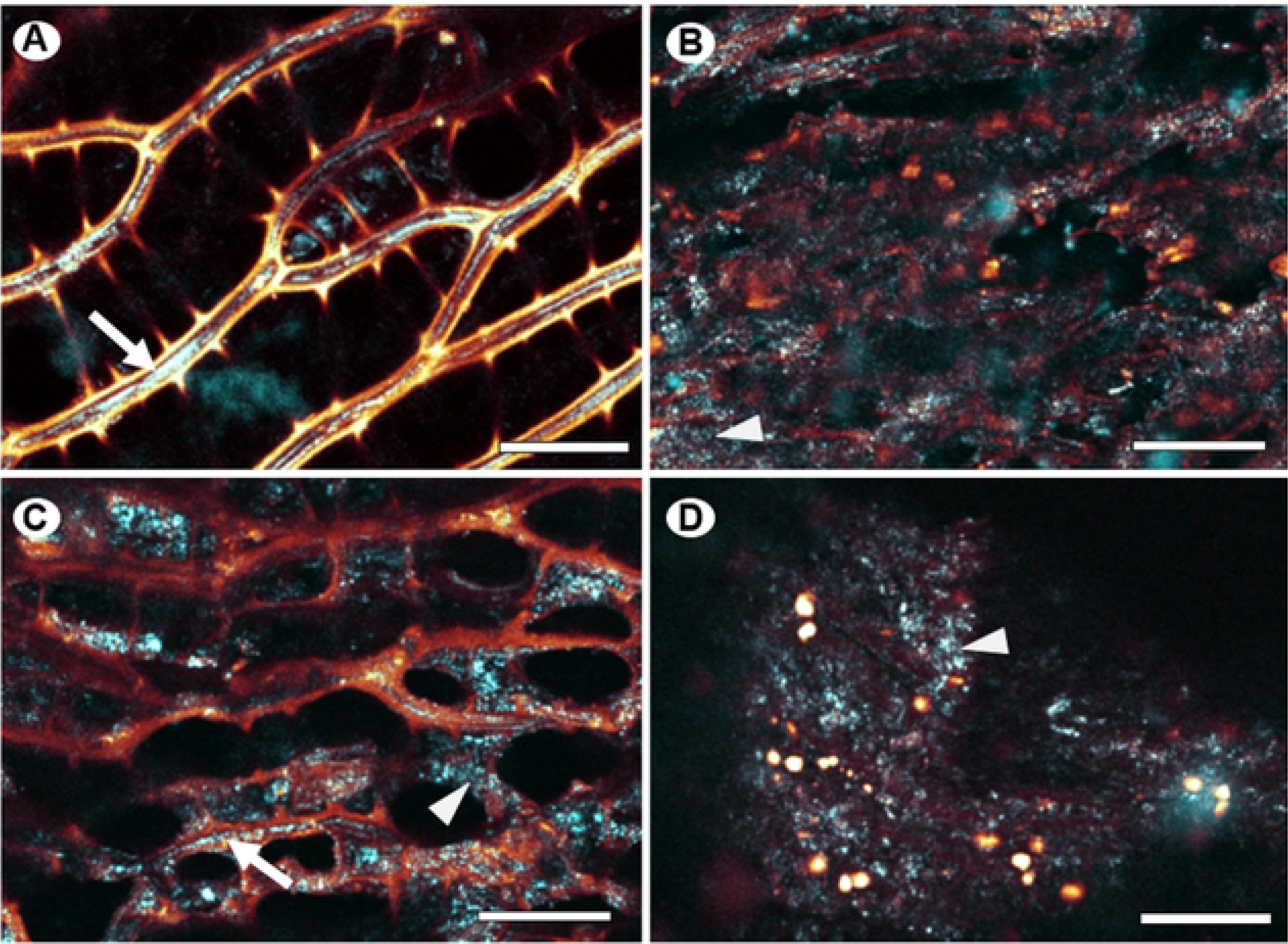
Confocal laser scanning microscopy of *Sphagnum fuscum* leaf material from different sources of peat moss. Acrotelm (A) and catotelm (B) collected in Abisko peat bog compared to peat (C) and peat pretreated at 180 °c(D). fl-1,4 glucans are visualized with calcofluor white staining and shown in cyan. The arrow points to intact parts of the primary cell wall within chlorophyllose cells, where presumably cellulose is clearly exposed to the dye. Fuzzy and speckled labeling may indicate degradation of calcofluor white epitopes (arrowheads). Autofluorescence from lignin-like compounds is shown in yellow/orange. The signal clearly localizes to the secondary cell walls of hyaline cells in A) and C), while it localizes to droplets in D). Scale bars, 25 µm.

The structure of hyaline and chlorophyllose cells in peat (Fig. 6C) resembled that of acrotelm, but the fluorescent signal from the lignin-like compounds was decreased compared to that of peat. Furthermore, the calcofluor white signal was less organized in peat, although a distinct line could be noted as thin lines in some places (Fig. 6C). Following the 180 °C pretreatment of peat (Fig. 6D), individual cell types could no longer be recognized and the calcofluor white signal was no longer localized to cell walls. The autofluorescence signal from lignin-like compounds was intensified and appeared to have relocated to droplets, similar to observations of wheat straw lignin relocation following pretreatment (Hansen et al., 2011). Together, these data indicate that a 180 °C pretreatment leads to extensive degradation of *Sphagnum* cell walls and lignin relocation.

## 4.0 Conclusions

Hydrothermal pretreatment of peat releases monosaccharides and glucose-rich oligosaccharides. These oligosaccharides are highly accessible to an advanced enzyme cocktail designed for saccharification of pretreated lignocellulosic material such as wheat straw. The insoluble part of the pretreated peat was saccharified at a somewhat higher overall rate compared to pretreated wheat straw. However, the high iron concentration in peat led to oxidative acidification of pretreated peat, which was alleviated by the addition of catalase. These oxidative reactions likely contributed to the rapid inactivation initially observed for the saccharifying enzyme cocktail.

Cellulose and lignin-like compounds in peat and catotelm were less well defined compared to their counterparts in the leaves of fresh *Sphagnum* moss. Hydrothermal pretreatment resulted in relocation of lignin-like components of the cell wall. The partial decomposition of polysaccharides and the relocation of lignin-like compounds induced by hydrothermal pretreatment result in a peat material that is just as susceptible to enzymatic saccharification as lignocellulose from vascular plants. These findings offer a new perspective on the recalcitrance of *Sphagnum* moss and the protection of carbon storage in peat bogs.

**Appendix A. Supplementary data**

**Appendix B. R-code**

## References

Angeltveit, C. F., Jeoh, T., & Horn, S. J. (2023). Lytic polysaccharide monooxygenase activity increases productive binding capacity of cellobiohydrolases on cellulose. Bioresource Technology, 389, 129806. 10.1016/j.biortech.2023.129806

Ballance, S., Borsheim, K., Inngjerdingen, K., Paulsen, B., & Christensen, B. (2007). A re-examination and partial characterisation of polysaccharides released by mild acid hydrolysis from the chlorite-treated leaves of Sphagnum papillosum. Carbohydrate Polymers, 67(1), 104–115. 10.1016/j.carbpol.2006.04.020

Benito-Román, Ó., Sanz, M. T., & Beltrán, S. (2024). Studies of degradation of pectin derived compounds from onion skins in subcritical water. The Journal of Supercritical Fluids, 206, 106155. 10.1016/j.supflu.2023.106155

Clymo, R. S. (1963). Ion Exchange in Sphagnum and its Relation to Bog Ecology. Annals of Botany, 27(2), 309–324. 10.1093/oxfordjournals.aob.a083847

Environment, U. N. (2024, February 28). Protecting peatlands for people and planet. UNEP -UN Environment Programme. http://www.unep.org/topics/ocean-seas-and-coasts/blue-ecosystems/protecting-peatlands-people-and-planet

Fabiola Rodríguez-Zúñiga, U., Cannella, D., Campos Giordano, R. de, Camargo Giordano, R. de L., Jørgensen, H., & Felby, C. (2015). Lignocellulose pretreatment technologies affect the level of enzymatic cellulose oxidation by LPMO. Green Chemistry, 17(5), 2896–2903. 10.1039/C4GC02179G

Freeman, C., Ostle, N., & Kang, H. (2001). An enzymic “latch” on a global carbon store. Nature, 409(6817), Article 6817. 10.1038/35051650

Glime, J. M. 2017. (n.d.). Scribd. Retrieved April 9, 2024, from https://www.scribd.com/document/708029110/Volume-1-Chapter-2-5-Bryophyta-Sphagnopsida

Hájek, T., Ballance, S., Limpens, J., Zijlstra, M., & Verhoeven, J. T. A. (2011). Cell-wall polysaccharides play an important role in decay resistance of Sphagnum and actively depressed decomposition in vitro. Biogeochemistry, 103(1), 45–57. 10.1007/s10533-010-9444-3

Hansen, M. A. T., Kristensen, J. B., Felby, C., & Jørgensen, H. (2011). Pretreatment and enzymatic hydrolysis of wheat straw (Triticum aestivum L.) – The impact of lignin relocation and plant tissues on enzymatic accessibility. Bioresource Technology, 102(3), 2804–2811. 10.1016/j.biortech.2010.10.030

Herbaut, M., Zoghlami, A., Habrant, A., Falourd, X., Foucat, L., Chabbert, B., & Paës, G. (2018). Multimodal analysis of pretreated biomass species highlights generic markers of lignocellulose recalcitrance. Biotechnology for Biofuels, 11(1), 52. 10.1186/s13068-018-1053-8

Ibbett, R., Gaddipati, S., Greetham, D., Hill, S., & Tucker, G. (2014). The kinetics of inhibitor production resulting from hydrothermal deconstruction of wheat straw studied using a pressurised microwave reactor. Biotechnology for Biofuels, 7(1), 45. 10.1186/1754-6834-7-45

Johansen, K. S. (2016). Discovery and industrial applications of lytic polysaccharide mono-oxygenases. Biochemical Society Transactions, 44(1), 143–149. 10.1042/BST20150204

Kabel, M. A., Bos, G., Zeevalking, J., Voragen, A. G. J., & Schols, H. A. (2007). Effect of pretreatment severity on xylan solubility and enzymatic breakdown of the remaining cellulose from wheat straw. Bioresource Technology, 98(10), 2034–2042. 10.1016/j.biortech.2006.08.006

Kremer, C., Pettolino, F., Bacic, A., & Drinnan, A. (2004). Distribution of cell wall components in Sphagnum hyaline cells and in liverwort and hornwort elaters. Planta, 219(6), 1023–1035.

Kristensen, J. B., Thygesen, L. G., Felby, C., Jorgensen, H., & Elder, T. (2008). Cell wall structural changes in wheat straw pretreated for bioethanol production. Biotechnology for Biofuels, 1(1), 5. 10.1186/1754-6834-1-5

Lehmann, J., & Kleber, M. (2015). The contentious nature of soil organic matter. Nature, 528(7580), 60–68. 10.1038/nature16069

Ligrone, R., Carafa, A., Duckett, J. G., Renzaglia, K. S., & Ruel, K. (2008). Immunocytochemical detection of lignin-related epitopes in cell walls in bryophytes and the charalean alga Nitella. Plant Systematics and Evolution, 270(3), 257–272. 10.1007/s00606-007-0617-z

Liu, C., Zhao, Y., Ma, L., Zhai, G., Li, X., Freeman, C., & Feng, X. (2024). Metallic protection of soil carbon: Divergent drainage effects in *Sphagnum* vs. non-*Sphagnum* wetlands. *National Science Review*, nwae178. 10.1093/nsr/nwae178

Merali, Z., Collins, S. R. A., Elliston, A., Wilson, D. R., Käsper, A., & Waldron, K. W. (2015). Characterization of cell wall components of wheat bran following hydrothermal pretreatment and fractionation. Biotechnology for Biofuels, 8(1), 23. 10.1186/s13068-015-0207-1

Miles, C. J., & Brezonik, P. L. (1981). Oxygen consumption in humic-colored waters by a photochemical ferrous-ferric catalytic cycle. Environmental Science & Technology, 15(9), 1089–1095. 10.1021/es00091a010

Moore, T., & Basiliko, N. (2006). Decomposition in Boreal Peatlands. In R. K. Wieder & D. H. Vitt (Eds.), Boreal Peatland Ecosystems (Vol. 188, pp. 125–143). Springer Berlin Heidelberg. 10.1007/978-3-540-31913-9_7

Nature restoration law—European Commission. Retrieved July 8, 2024, from https://environment.ec.europa.eu/publications/nature-restoration-law_en

Ni, B., Yu, X., Duan, X., & Zou, Y. (2024). Wetland soil organic carbon balance is reversed by old carbon and iron oxide additions. Frontiers in Microbiology, 14. 10.3389/fmicb.2023.1327265

Osborne, C., Gilbert-Parkes, S., Spiers, G., Lamit, L. J., Lilleskov, E. A., Basiliko, N., Watmough, S., Andersen, R., Artz, R. E., Benscoter, B. W., Bragazza, L., Bräuer, S. L., Carson, M. A., Chen, X., Chimner, R. A., Clarkson, B. R., Enriquez, A. S., Grover, S. P., Harris, L. I., … Global Peatland Microbiome Project. (2024). Global Patterns of Metal and Other Element Enrichment in Bog and Fen Peatlands. Archives of Environmental Contamination and Toxicology, 86(2), 125–139. 10.1007/s00244-024-01051-3

Page, S. E., Kling, G. W., Sander, M., Harrold, K. H., Logan, J. R., McNeill, K., & Cory, R. M. (2013). Dark Formation of Hydroxyl Radical in Arctic Soil and Surface Waters. Environmental Science & Technology, 47(22), 12860–12867. 10.1021/es4033265

Painter, T. J. (1983). Residues of d-*lyxo*-5-hexosulopyranuronic acid in *Sphagnum* holocellulose, and their role in cross-linking. Carbohydrate Research, 124(1), C18–C21. 10.1016/0008-6215(83)88373-6

Painter, T. J. (1991). Lindow man, tollund man and other peat-bog bodies: The preservative and antimicrobial action of Sphagnan, a reactive glycuronoglycan with tanning and sequestering properties. Carbohydrate Polymers, 15(2), 123–142. 10.1016/0144-8617(91)90028-B

Peciulyte, A., Samuelsson, L., Olsson, L., McFarland, K. C., Frickmann, J., Østergård, L., Halvorsen, R., Scott, B. R., & Johansen, K. S. (2018). Redox processes acidify and decarboxylate steam-pretreated lignocellulosic biomass and are modulated by LPMO and catalase. Biotechnology for Biofuels, 11(1), 165. 10.1186/s13068-018-1159-z

Petersen, M. Ø., Larsen, J., & Thomsen, M. H. (2009). Optimization of hydrothermal pretreatment of wheat straw for production of bioethanol at low water consumption without addition of chemicals. Biomass and Bioenergy, 33(5), 834–840. 10.1016/j.biombioe.2009.01.004

Pińkowska, H., Krzywonos, M., Wolak, P., & Złocińska, A. (2019). Production of uronic acids by hydrothermolysis of pectin as a model substance for plant biomass waste. Green Processing and Synthesis, 8(1), 683–690. 10.1515/gps-2019-0039

Pipes, G. T., & Yavitt, J. B. (2022). Biochemical components of Sphagnum and persistence in peat soil. Canadian Journal of Soil Science, 102(3), 785–795. 10.1139/cjss-2021-0137

R Core Team (2024). R: A Language and Environment for Statistical Computing. R Foundation for Statistical Computing, Vienna, Austria. <https://www.R-project.org/>.

Riedel, T., Zak, D., Biester, H., & Dittmar, T. (2013). Iron traps terrestrially derived dissolved organic matter at redox interfaces. Proceedings of the National Academy of Sciences, 110(25), 10101–10105. 10.1073/pnas.1221487110

Roberts, A. W., Lahnstein, J., Hsieh, Y. S. Y., Xing, X., Yap, K., Chaves, A. M., Scavuzzo-Duggan, T. R., Dimitroff, G., Lonsdale, A., Roberts, E., Bulone, V., Fincher, G. B., Doblin, M. S., Bacic, A., & Burton, R. A. (2018). Functional Characterization of a Glycosyltransferase from the Moss Physcomitrella patens Involved in the Biosynthesis of a Novel Cell Wall Arabinoglucan. The Plant Cell, 30(6), 1293–1308. 10.1105/tpc.18.00082

Romanowicz, K. J., Kane, E. S., Potvin, L. R., Daniels, A. L., Kolka, R. K., & Lilleskov, E. A. (2015). Understanding drivers of peatland extracellular enzyme activity in the PEATcosm experiment: Mixed evidence for enzymic latch hypothesis. Plant and Soil, 397(1), 371–386. 10.1007/s11104-015-2746-4

Santana, M. A. E., & Okino, E. Y. A. (2007). Chemical composition of 36 Brazilian Amazon forest wood species. 61(5), 469–477. 10.1515/HF.2007.084

Scott, B. R., Huang, H. Z., Frickman, J., Halvorsen, R., & Johansen, K. S. (2016). Catalase improves saccharification of lignocellulose by reducing lytic polysaccharide monooxygenase-associated enzyme inactivation. Biotechnology Letters, 38(3), 425–434. 10.1007/s10529-015-1989-8

Selwyn, M. J. (1965). A simple test for inactivation of an enzyme during assay. Biochimica et Biophysica Acta (BBA) - Enzymology and Biological Oxidation, 105(1), 193–195. 10.1016/S0926-6593(65)80190-4

Shotyk, W. (1988). Review of the inorganic geochemistry of peats and peatland waters. Earth-Science Reviews, 25(2), 95–176. 10.1016/0012-8252(88)90067-0

Sidiras, D., Batzias, F., Ranjan, R., & Tsapatsis, M. (2011). Simulation and optimization of batch autohydrolysis of wheat straw to monosaccharides and oligosaccharides. Bioresource Technology, 102(22), 10486–10492. 10.1016/j.biortech.2011.08.059

Soetaert, K., Petzoldt, T., & Setzer, R. W. (2010). Solving Differential Equations in R: Package deSolve. Journal of Statistical Software, 33, 1–25. 10.18637/jss.v033.i09

Sun, S., Wen, J., Sun, S., & Sun, R.-C. (2015). Systematic evaluation of the degraded products evolved from the hydrothermal pretreatment of sweet sorghum stems. Biotechnology for Biofuels, 8(1), 37. 10.1186/s13068-015-0223-1

Thormann, M. N., Currah, R. S., & Bayley, S. E. (2002). The relative ability of fungi from Sphagnum fuscum to decompose selected carbon substrates. Canadian Journal of Microbiology, 48(3), 204–211. 10.1139/w02-010

Thygesen, A., Thomsen, M. H., Jørgensen, H., Christensen, B. H., & Thomsen, A. B. (2004, May). Hydrothermal treatment of wheat straw on pilot plant scale. In 2nd World Conference and Technology Exhibition on Biomass for Energy, Industry and Climate Protection, Rome, Italy.

Trusiak, A., Treibergs, L. A., Kling, G. W., & Cory, R. M. (2018). The role of iron and reactive oxygen species in the production of CO2 in arctic soil waters. Geochimica et Cosmochimica Acta, 224, 80–95. 10.1016/j.gca.2017.12.022

Tsuneda, A., Thormann, M. N., & Currah, R. S. (2001). Modes of cell-wall degradation of *Sphagnum fuscum* by *Acremonium* cf. *Curvulum* and *Oidiodendron maius*. Canadian Journal of Botany, 79(1), 93–100. 10.1139/b00-149

Urbanová, Z., & Hájek, T. (2021). Revisiting the concept of ‘enzymic latch’ on carbon in peatlands. Science of The Total Environment, 779, 146384. 10.1016/j.scitotenv.2021.146384

van Breemen, N. (1995). How *Sphagnum* bogs down other plants. Trends in Ecology & Evolution, 10(7), 270–275. 10.1016/0169-5347(95)90007-1

Verhoeven, J. T. A., & Liefveld, W. M. (1997). The ecological significance of organochemical compounds in Sphagnum. Acta Botanica Neerlandica, 46(2), 117–130.

Wang, Y., Wang, H., He, J.-S., & Feng, X. (2017). Iron-mediated soil carbon response to water-table decline in an alpine wetland. Nature Communications, 8(1), 15972. 10.1038/ncomms15972

Wang, Z.-W., Zhu, M.-Q., Li, M.-F., Wang, J.-Q., Wei, Q., & Sun, R.-C. (2016). Comprehensive evaluation of the liquid fraction during the hydrothermal treatment of rapeseed straw. Biotechnology for Biofuels, 9(1), 142. 10.1186/s13068-016-0552-8

Wen, Y., Zang, H., Ma, Q., Evans, C. D., Chadwick, D. R., & Jones, D. L. (2019). Is the ‘enzyme latch’ or ‘iron gate’ the key to protecting soil organic carbon in peatlands? Geoderma, 349, 107–113. 10.1016/j.geoderma.2019.04.023

Weng, J.-K., & Chapple, C. (2010). The origin and evolution of lignin biosynthesis. New Phytologist, 187(2), 273–285. 10.1111/j.1469-8137.2010.03327.x

Westereng, B., Agger, J. W., Horn, S. J., Vaaje-Kolstad, G., Aachmann, F. L., Stenstrøm, Y. H., & Eijsink, V. G. H. (2013). Efficient separation of oxidized cello-oligosaccharides generated by cellulose degrading lytic polysaccharide monooxygenases. Journal of Chromatography. A, 1271(1), 144–152. 10.1016/j.chroma.2012.11.048

Wilson, R., Hopple, A., Tfaily, M., Sebestyen, S., Schadt, C., Pfeifer-Meister, L., Zalman, C., Mcfarlane, K., Kostka, J., Kolton, M., Kolka, R., Kluber, L., Keller, J., Guilderson, T., Griffiths, N., Chanton, J., Bridgham, S., & Hanson, P. (2016). Stability of Peatland Carbon to Rising Temperatures. Nature Communications, 7. 10.1038/ncomms13723

Wolfenden, R., & Snider, M. J. (2001). The Depth of Chemical Time and the Power of Enzymes as Catalysts. Accounts of Chemical Research, 34(12), 938–945. 10.1021/ar000058i

Xu, C., Zhang, X., Hussein, Z., Wang, P., Chen, R., Yuan, Q., Gao, Y., Song, N., & Gouda, S. G. (2021). Influence of the structure and properties of lignocellulose on the physicochemical characteristics of lignocellulose-based residues used as an environmentally friendly substrate. Science of The Total Environment, 790, 148089. 10.1016/j.scitotenv.2021.148089

Ye, Z.-H., & Zhong, R. (2022). Cell wall biology of the moss *Physcomitrium patens*. Journal of Experimental Botany, 73(13), 4440–4453. 10.1093/jxb/erac122

Yu, Z. C. (2012). Northern peatland carbon stocks and dynamics: A review. Biogeosciences, 9(10), 4071–4085. 10.5194/bg-9-4071-2012

Zakaria, M. R., Hirata, S., & Hassan, M. A. (2015). Hydrothermal pretreatment enhanced enzymatic hydrolysis and glucose production from oil palm biomass. Bioresource Technology, 176, 142–148. 10.1016/j.biortech.2014.11.027

Zhao, Y., Liu, C., Li, X., Ma, L., Zhai, G., & Feng, X. (2023). Sphagnum increases soil’s sequestration capacity of mineral-associated organic carbon via activating metal oxides. Nature Communications, 14(1), 5052. 10.1038/s41467-023-40863-0

